# Quantitative evaluation of two-photon calcium imaging modalities for high-speed volumetric calcium imaging in scattering brain tissue

**DOI:** 10.1101/115659

**Authors:** Siegfried Weisenburger, Robert Prevedel, Alipasha Vaziri

## Abstract

Considerable efforts are currently being devoted to enhance the speed, spatial resolution and the size of the 3D sample volumes in which calcium imaging methods can capture neuronal network activity in different model systems. In the mammalian brain, tissue scattering severely limits the use of parallel acquisition techniques such as wide-field imaging and, as a consequence, methods based on two-photon point-scanning (2PM) have become the method of choice. However, 2PM faces severe restrictions due to technical limitations such as scan speed, laser power, and those related to the fluorescent probes, calling for conceptually new approaches to enhance the performance of two-photon calcium imaging schemes. Here we provide a detailed quantitative evaluation and comparison of different excitation/detection modalities from the perspective of detecting neuronal activity that are based on different point-spread functions (PSF), laser repetition rates and sampling strategies. We demonstrate the conditions for which imaging speed and signal-to-noise ratio are optimized for a given average power. Our results are based on numerical simulations which are informed by experimentally measured parameters and show that volumetric field of view and acquisition speed can be considerably improved compared to traditional 2PM schemes by a holistic optimization approach.

## Introduction

Unraveling the principles of how neuronal networks process and represent sensory information and how these ongoing dynamics produce behavioral output is at the forefront of current neuroscience research [1-5]. To address these questions, imaging techniques are required that have the ability to record the neuronal activity from large networks with high temporal and single-cell spatial resolution. Calcium (Ca^2+^) imaging using genetically encoded Ca^2+^ indicators (GECI) such as GCaMP offers a relatively noninvasive approach to read out neuronal activity optically from genetically defined populations [6-10]. The development of these fluorescent labels has subsequently fueled the inception of complementary fast, high resolution and large-scale optical microscopy methods for recording of neuronal activity in various model organisms, including the recent demonstrations of whole-brain Ca^2+^ imaging in small model systems [11-14].

In semi-transparent and small model organisms such as *C. elegans* and zebrafish larvae, one-photon excitation combined with wide-field detection schemes has allowed the realization of various types of high-speed Ca^2+^ imaging methodologies, such as light-sheet microscopy modalities [13, 15, 16, 17], light-field microscopy [14, 18, 19], or methods based on multi-plane imaging [20-23]. While these approaches have in common that their underlying parallelization of acquisition through wide-field excitation and a 2D-detector-array-based detection leads to a significant neuronal sampling rate, the performance of these wide-field detection schemes degrades severely with increasing imaging depth due to scatter induced pixel cross talk, which restricts their applicability to thin or weakly scattering specimens.

In scattering tissue such as the mammalian brain, excitation using two-photon absorption in combination with point-scanning microscopy has become the gold standard for recording neuronal activity as it provides the necessary lateral and axial resolution, signal-to-background ratio (SBR) and improved depth penetration in biological tissue [24]. However, conventional two-photon scanning microscopy (2PM), which uses a diffraction-limited excitation spot, typically struggles to achieve sufficient temporal resolution to resolve Ca^2+^ dynamics and to faithfully sample all active neurons within a large neuronal population. This is despite of the various efforts to improve the performance of 2PM by using remote axial scanning [25-28], acoustooptical scanners [29-31] or multiplexing [22, 23, 32-35]. In addition to all technical hardware limitations, the ultimate limitation in the obtainable sample rate in all diffraction-limited scanning approaches is given by the properties of the fluorophores such as emission rate saturation due to fluorescence lifetime and photodamage, which put serious constrains on the obtainable recording speed.

For the most part, a state-of-the-art 2P scanning based Ca^2+^ imaging microscope achieves frame rates of ~30 Hz for a 500×500 μm field of view (FOV) (512×512 pixels) in the case of bidirectional scanning with a resonant scanner [36, 37]. This video-rate temporal resolution is adequate to record fast Ca^2+^ dynamics in 2D, but scanning a 3D volumetric FOV (V-FOV) covering 500×500×500 μm (512×512×500 pixels) and assuming 500 axial planes scanned with e.g. a piezo would yield a volume rate of only ~0.06 Hz, clearly insufficient to faithfully sample neuronal dynamics for each neuron within such a volume. In order to sample the same volume at somewhat more physiologically relevant time scales (e.g. >5 Hz) would put extreme demands on the necessary laser technology, scan and data acquisition hardware but most importantly would be severely limited by the photo-physics of fluorescence itself: To sample the above volume consisting of ^~^ 0.5×10^9^ voxels at 5 Hz with a diffraction-limited 2PM would require a laser source with >2.5 GHz repetition rate. If a laser with that repetition rate existed, at tolerable average powers on the order of 250 mW [38, 39], the excitation pulse energy would be around 100 pJ, likely insufficient to yield sufficient fluorescence signals. Finally, at such high repetition rates, the dwell times of < 0.3 ns would be an order of magnitude shorter than the fluorescence lifetime of the typically used fluorophores, thus leading to pixel cross talk and signal degradation.

Addressing these challenges requires conceptually new imaging approaches. In recent work, we proposed and experimentally demonstrated that spatial resolution can be traded in favor of V-FOV, imaging speed and fluorescence signal [40]. Since the typical size of a mouse cortical neuron is on the order 10-20 μm [41], we reasoned that an enlarged, isotropic PSF can be utilized for scanning while maintaining single-cell resolution. Due to the larger extent of the PSF, the volume can be sampled with the minimally required number of excitation voxels. At the same time, this approach maximizes the signal and signal-to-noise ratio (SNR) of the recordings that can be obtained from Ca^2+^ indicator-expressing neurons in the mouse brain. In the present work, we numerically evaluate and compare this conceptually new imaging approach which uses an enlarged, sculpted excitation volume to other, more conventional two-photon imaging modalities that are based on diffraction-limited PSF.

Our work is based on numerical simulations which are informed by experimentally measured parameters, thus representing practical and real-world imaging conditions. We show that this new approach leads to an optimal trade-off between signal, signal-to-noise ratio, imaging speed, resolution and V-FOV, and is well-suited for systems neuroscience applications where a reduced spatial resolution is tolerable for the benefit of a highly improved volumetric FOV and acquisition speed. We discuss the results with regards to important input parameters as well as their mutual trade-offs, and show that an excitation scheme in which a single laser pulse per image voxel is chosen leads to the optimal imaging performance. We further show that the image quality and reduced spatial resolution is indeed sufficient for faithful identification and segmentation of neurons and extraction of Ca^2+^ dynamics using cell segmentation and signal demixing approaches.

## Enlarged, sculpted PSF improves Ca^2+^-imaging performance

A common goal in 2PM is to increase the size of the acquisition volume, i.e. V-FOV, while maintaining high temporal resolution and sensitivity, i.e. signal-to-noise ratio. Achieving these goals simultaneously has been notoriously difficult because of the inverse relationship between the size of the total scanned volume and the signal collected per voxel within the voxel dwell time. In principle, faster scanners would allow for faster frame rates, if excitation power could be increased appropriately to compensate for the shorter voxel dwell time. However, fluorophore saturation as well as side effects from high excitation intensities such as heating, photo-damage and photo-bleaching ultimately limit the maximally possible excitation and emission rate even in the case of unlimited laser power. In light of these practical constraints, our study aimed at comparing excitation and detection schemes to optimize the imaging performance for recording neuronal dynamics with two-photon microscopes and current Ca^2+^ indicators.

Common point-scanning microscopes based on two-photon excitation aim to produce diffraction-limited PSFs in the sample. This is done primarily in order to achieve good spatial resolution, especially in the axial direction, so that the excitation is confined to a small volume, typically on the order of 1 μm^3^. However, for many questions in system neuroscience such a high spatial resolution may not be necessary and would be rather traded off by the experimenter with a larger volume size or higher speed. An important quantity in current neuroscience is the ‘recording capacity’, i.e. the number of neurons whose activity can be read-out near-simultaneously. This requires single-cell spatial resolution at an adequate signal-to-noise (SNR) for automatic cell detection during the analysis and with sufficiently high temporal resolution to follow activity-evoked Ca^2+^ transients. More concretely, to effectively sample neuronal soma in rodent brains whose average size is >10 μm, a spatial resolution of ~5×5×5 μm should be adequate according to the sampling theorem. At the same time, a temporal resolution of >5 Hz is sufficient to record Ca^2+^-evoked dynamics from indicators such as GCaMP, which show typical Ca^2+^ transients with decay times of ~0.3 − 1 s upon stimulation.

Unfortunately, isotropic PSFs extending several micrometers in size are not straightforward to realize because of the intrinsic, non-linear relationship between the lateral (*w*_0_) and axial size (*z*) of a Gaussian focus, *z ~ w_0_^2^.* Nonetheless, strategies from ultra-fast optics exist that enable arbitrary shaping of focal volumes. Such light-sculpting approaches are possible, for example by using the technique of temporal focusing (TeFo) [11, 40, 42-45], which allows an effective decoupling of the axial and lateral confinement of the excitation volume.

Adjusting the PSF to the object of interest results in three significant advantages: (*i*) It reduces the number of voxels that need to be scanned per unit volume to the theoretical minimum that is necessary to resolve a structure of interest (e.g. neuronal cell bodies). This in turn implies faster frame or volume acquisition rates or larger (volumetric) FOV at the same temporal resolution. (*ii*) Exciting less voxels requires a lower minimal laser pulse repetition rate, thus permitting higher pulse energies and hence higher fluorescence signal for the same average laser power. In addition, the lower repetition rate and correspondingly longer signal acquisition intervals of the laser mean that all fluorescence emitted within the fluorescence life time of the fluorophore can be collected. (*iii*) The overall signal-to-noise ratio per voxel at a given excitation intensity is increased compared to the single diffraction-limited PSF. Since all fluorophores are excited with a single pulse, the signal benefits nonlinearly. Consequently, less intensity (average power per unit area) is required to generate a given SNR and thus illumination can remain below fluorophore saturation and nonlinear photo-damage thresholds.

In the following, we will elaborate on the above stated advantages of using a light-sculpted (LS) PSF. The size of the point-spread function, *A*_PSF_, influences the acquisition speed for a given FOV as follows: The plane exposure time *t*_exp_ necessary to acquire the signal from the desired field-of-view, *A*_FOV_, depends on the excitation area *A*_PSF_ and its dwell time Δ*t*, via *t*_exp_ = (*A*_FOV_ / *A*_PSF_) × Δ*t*. For a constant voxel dwell time, therefore the largest possible value for *A*_PSF_ for a desired spatial resolution will result in the minimal number of voxels that is required to be scanned and dwelled over for time *Δt* and thus will result in the shortest plane exposure time *t*_exp_.

The actual frame or volume rate depends on the dwell time, which naturally needs to be sufficiently long so that enough fluorescence signal can be acquired. The fluorescence signal in two-photon excitation via a pulsed laser source is proportional to the number of absorbed photons per excitation volume per pulse, *N_a_*, and is given by [46]:

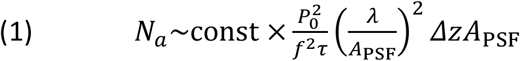

with *P*_0_ denoting the average laser power at the sample plane, *f* the laser’s pulse repetition rate, *τ* the pulse length, *λ* the central wavelength, *A*_PSF_ the excitation area, and Δ*z* the axial excitation confinement at the sample. The constant is further const ∝ *σ_2_*_PA_ × *C,* where *σ_2_*_PA_ is the 2PA cross section (in units of GM = 10^− 50^ cm^4^ s photon^− 1^) of the fluorophore, and *C* the concentration of the fluorophores in the excitation volume which determines the number of active fluorophores.

Taking into account the finite voxel dwell time Δ*t*, the number of absorbed photons per excitation volume then becomes

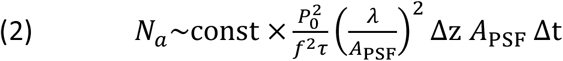

It is immediately obvious that the total fluorescence signal from each voxel scales with the pulse energy squared, i.e. *N_a_~*(*P*_0_*/f*)^2^, therefore the lowest possible laser repetition rate for a given average power would result in the highest fluorescence signal. Taking into account that each imaging voxel needs at least one pulse for excitation, we conclude that the fluorescence signal is maximized when one pulse per pixel is used. Furthermore, Equation (2) allows us to directly compare the signal obtained with excitation PSFs of different sizes (characterized by their volume *V* = *Δz* A_PSF_). In our particular case, we will contrast the effects on obtainable signal and required laser power (and intensity) when using an isotropic, enlarged PSF compared to a more conventionally used diffraction-limited PSF. Assuming an enlarged, light-sculpted (LS) PSF of *A* = 5×5 μm and Δz = 5 μm, in contrast to a diffraction-limited (DL) PSF of ^~^*A* = 0.5×0.5 μm and Δz = 1 μm, we can make the following illustrative calculations: At the same excitation intensity, (i.e. power density or average power per unit area, *I* = P_0_/A), the resulting signal will be a factor *Q* = *A*_LS_ Az_LS_ / A_DL_ Az_DL_ = 500x higher in the case of using a light-sculpted PSF. Correspondingly, to obtain the same fluorescence signal from a single voxel requires a power density of only 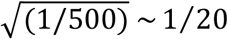 compared to the diffraction-limited case.

In practice, however, comparing the expected performance of our LS-PSF scheme of an enlarged, sculpted PSF together with a one-pulse-per-voxel acquisition scheme to the conventional, diffraction-limited PSF is slightly more complex. The reason for this is that compared to a configuration in which each neuron is excited by only a few (2-3) excitation spots, a diffraction limited scanning modality results in a larger number of voxels on a given neuron whose averaged signal contribute to the overall signal. However, in order to maintain the same acquisition rate in the diffraction limited configuration as in the LS-PSF configuration the voxel dwell time must be reduced to 1/V, and the pulse repetition rate correspondingly increased by V. This results in a decrease of pulse energy and thus a decrease of fluorescence signal. Overall, assuming the same power density (*P*_0_*/A*) the average (ave) signal from the sum of all *V* voxels that comprise the same volume as the sculpted PSF:

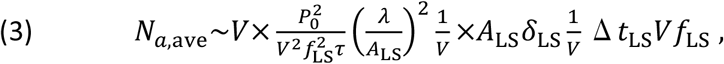

which overall is by a factor of *V*^2^ lower than the enlarged PSF signal. This non-linear, ~*V*^2^ increase in fluorescence signal for the enlarged PSF stems from the fact that it reduces the overall number of voxels to be scanned, which results in faster acquisition rates as well as in a correspondingly lower demand in laser repetition rate, which in turn leads to higher pulse energies and thus higher fluorescence signal.

Ultimately, every acquired signal needs to be resolved on top of noise, which can stem from various sources such as photon shot noise, fluorophore fluctuations and various electronic noise sources related to the data acquisition. The ultimately unavoidable noise is shot noise, which is identical in both excitation schemes. We note however that other types of noise such as the electronic noise or read-out noise can accumulate during the acquisition in the case of the diffraction-limited excitation, where several signals are read-out and integrated for each neuron.

Since photon shot noise scales with the square root of the signal, the signal-to-noise ratio, SNR, becomes 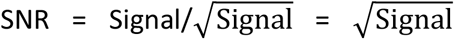. We thus find that 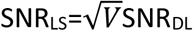 when compared to the single diffraction-limited PSF or the sparse sampled 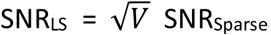, and SNRLS = *V* SNR_ave_ for multiple averaged diffraction-limited excitation. Thus, the signal-to-noise ratio improves at least by a factor 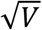. This effective gain in SNR can in turn be used to lower the excitation power density compared to conventional laser scanning microscopy, in order to reduce photo-damage and photo-bleaching, and to circumvent limitations due to the saturation of the fluorophores. Alternatively, for the same power density this gain can be used to lower the dwell time of each voxel in order to speed up image acquisition.

## In-silico comparison of imaging approaches

In order to quantitatively study and compare the performances of standard 2PM schemes and our LS-PSF method, we performed numerical simulations employing realistic experimentally obtained imaging parameters and fluorophore properties.

For our numerical simulations, we chose the following imaging parameters, as we regard them as most relevant for current open questions in system neuroscience:

- Field-of-view (FOV): 500×500×500 μm
- Fluorophore: GCaMP6m (σ = 20 GM, τ = 0.7 sec) (Ref. [6])
- Fluorophore concentration: approx. 10 μM (Ref. [47])
- Neuron size: 10 − 20 μm diameter (Ref. [41])
- Neuron density: 10^5^/mm^3^ (Ref. [41, 48, 49])
- Saturation/Photodamage: 20 nJ/μm^2^ (Ref. [50-52])
- Brain heating limit: 250 mW (Ref. [39])
- Laser pulse duration: 150 fs
- Laser wavelength: 1 μm
- Laser repetition rate: equal to number of voxels to be scanned per second (one-pulse-per voxel scheme)

For ease of discussion, we have assumed these parameters as fixed, since we do not consider the development of improved Ca^2+^ indicators but rather focus on the optimization of the fluorophore signal based on different imaging modalities. Furthermore, we do not consider any technical limitation due to hardware performance such as available laser power, energy, and mechanical scan speed.

In our simulations, we focused on the following four, conceptually different excitation modalities for our quantitative comparison study (Fig. 1):

a. Light-sculpted PSF (**LS-PSF**) with a spot size of 5×5×5 μm.
b. Diffraction-limited PSF with sparse sampling (**sparse-PSF**) and spot size 1×1×1 μm, but sampled only every 5×5×5 μm to produce the same number of pixels as in Fig. 1a.
c. Diffraction-limited PSF (**ave-PSF**) with spot size 1x1x1 μm, but signal binned into pixels of size 5×5×5 μm to produce the same number of pixels as in Fig. 1a.
d. Conventional diffraction-limited PSF (**DL-PSF**) with spot size 1×1×1 μm.

**Fig. 1:**
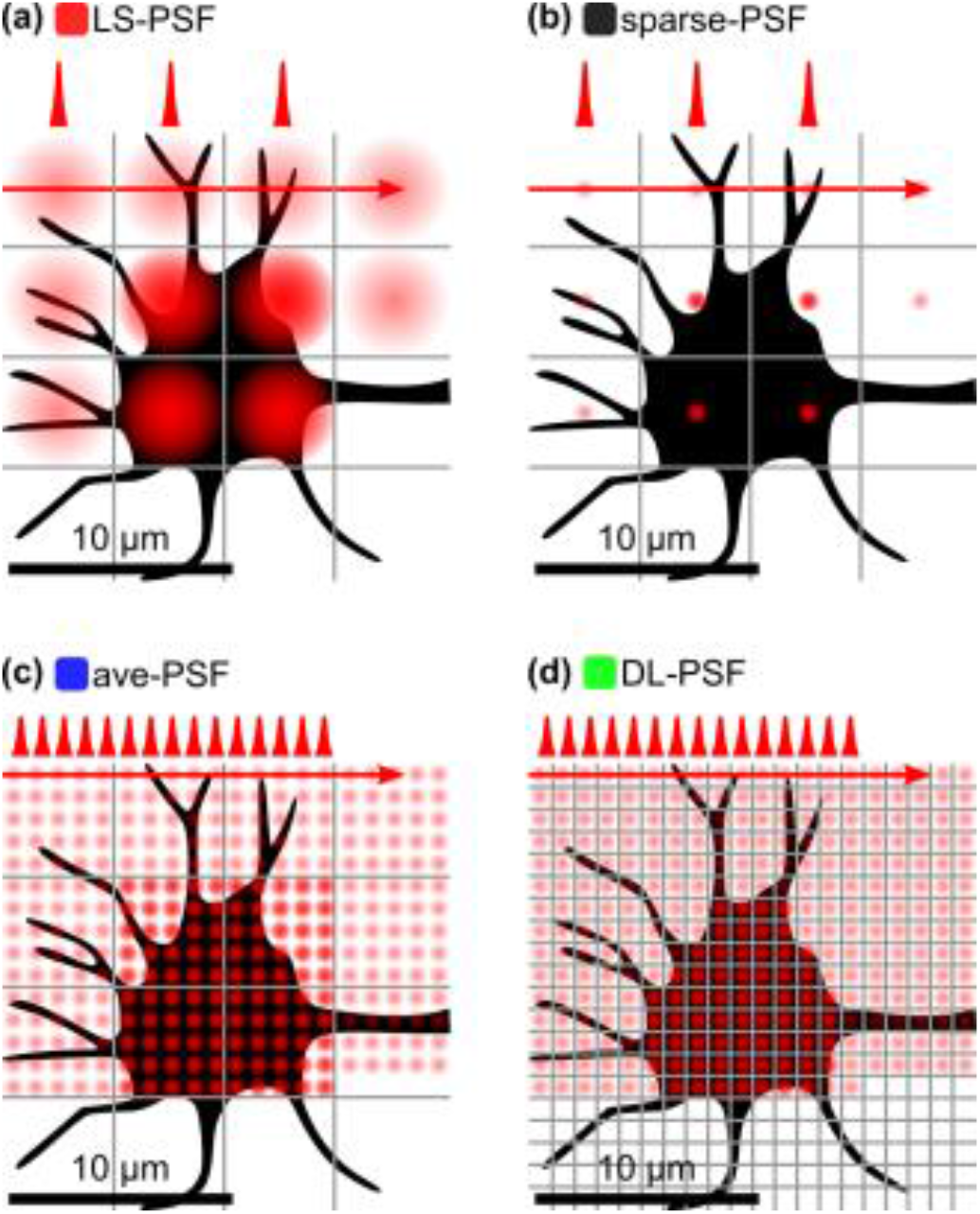
Imaging modalities considered in the simulations. (a) Light-sculpted, LS-PSF with a spot size of 5x5x5 μm, (b) diffraction-limited PSF (1×1×1 μm) with sparse sampling to produce the same number of pixels as in panel a, (c) diffraction-limited PSF with signal binned to produce the same number of pixels as in panel a, (d) Conventional diffraction-limited PSF.

The output of our simulations is two-dimensional pseudo-raw images (see Fig. 2 for the ground truth (a) and an example image (b)), whose pixel size matches the resolution of the imaging modality. The ‘intensity’ of each pixel is calculated from first principles following Eq. 1, and converted to pixel counts assuming a PMT sensitivity of 200 μW/A with 10^4^ gain, a PMT quantum efficiency of 40%, a preamplifier gain of 10^5^, and the signal connected into a 1 M*Ω* load. The results of the simulation in terms of signal intensity were cross-checked and validated against actual experimental measurements in our lab acquired in the LS-PSF (with the s-TeFo microscope from Ref. [40]) and DL-PSF (Scientifica 2-photon scanning microscope) scheme in the *in-vivo* mouse, expressing cytoplasmic GCaMP6m in the mouse motor cortex.

**Fig. 2:**
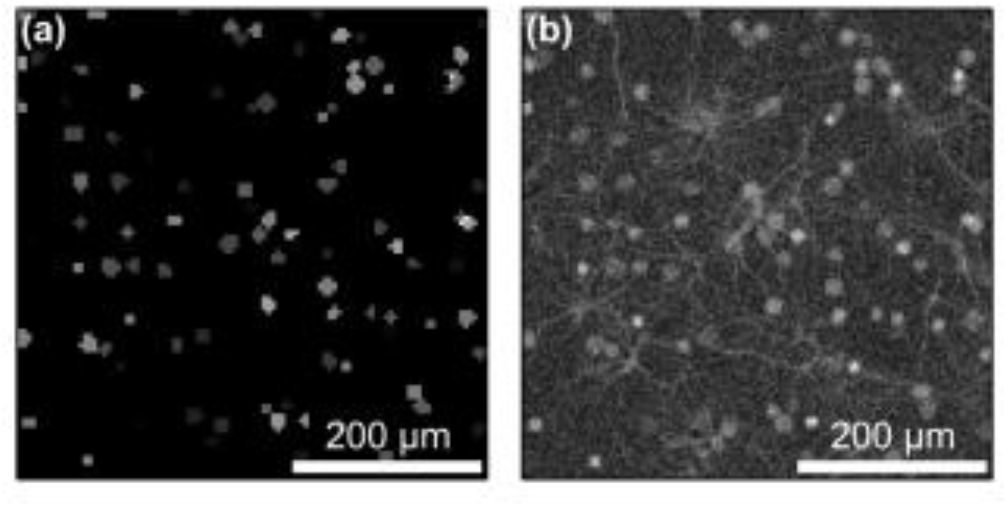
Comparison of simulated images and experiment. (a) Ground truth for the simulation without neuropil, showing the location and size of neurons on the 5×5 μm grid. (b) Simulated image with neurons and neuropil (axons, dendrites, etc). Example for the case of a diffraction-limited PSF.

The simulated images and their temporal dynamics were generated as follows: Neurons of varying diameter (10 − 20 μm) were randomly placed inside the imaging FOV at the average neuron density in the cortex, and their Ca^2+^ dynamics were modeled in time by assigning random Ca^2^+ dynamic kernels [10] with a decay time of 1s, random Ca^2^+ intensity changes between 20 and 100 % DF/F per transient, and a density of 0.1 transients/s. Background fluorescence was modeled by an image offset with normally distributed noise (values were determined from histograms of experimentally acquired images and scaled). For all signals and background contributions, shot noise was taken into account. Furthermore, background fluorescence stemming from neuropil such as dendrites, axons, etc., has been modeled by overlaying an artificial image of neuropil with their Ca^2+^ dynamics stemming from the mixed signal of ten randomly chosen neurons (values for the intensity scaling were determined from experimentally acquired images). The latter is important in the context of this study, since neuropil signal has to be properly detected and distinguished from neuronal (i.e. cell body) Ca^2+^ signals.

## Main results

### Signal-to-noise performance of various imaging modalities

To quantitatively compare the performance of the various imaging modalities, we simulated and analyzed the pseudo-images in respect to the following parameters and measures. Figure 3 shows the image signal-to-noise ratio (SNR), defined as

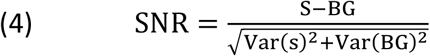

with S and BG denoting the signal and background values, respectively, depending on the power density incident on the sample for different voxel dwell times and hence volume recording rates.

**Fig. 3:**
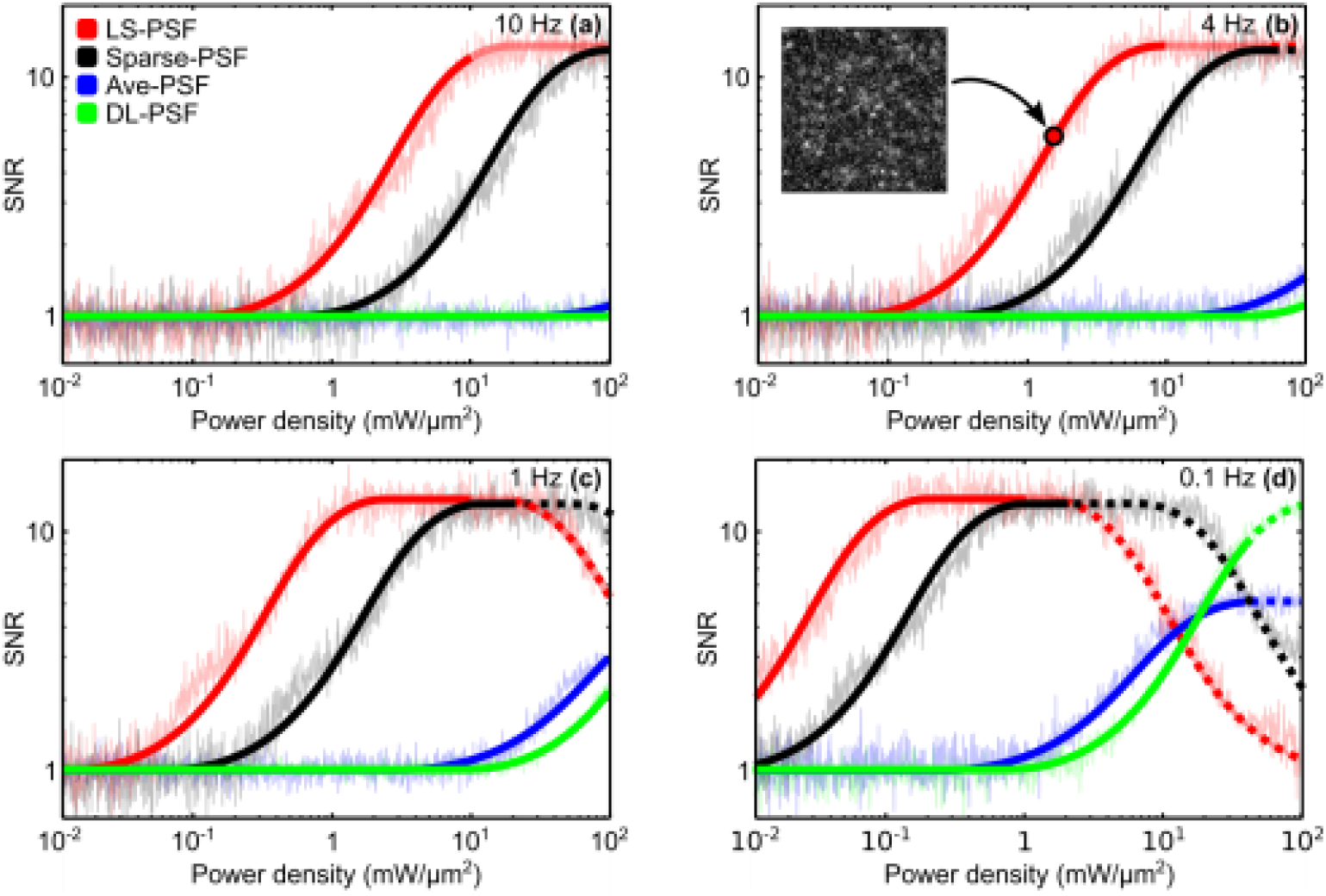
Signal-to-noise ratio as a function of incident power density for the different imaging modalities LS-PSF (light sculpted), sparse-PSF (diffraction-limited PSF with sparse scanning), ave-PSF (diffraction-limited PSF and averaging), and DL-PSF (diffraction-limited). The SNR is plotted for (a) 10 Hz volume recording rate (corresponding to 100 ns dwell time), (b) 4 Hz volume recording rate (250 ns dwell time), (c) 1 Hz volume recording rate (1 μs dwell time) and (d) 0.1 Hz volume recording rate (10 μs dwell time). The inset shows a corresponding simulated image for a selected data point. Semitransparent lines indicate that the total average power exceeds 250 mW, indicating limitation due to potential heating. Dashed lines mark pulse energy densities above the reported onset of non-linear photodamage of 20 nJ/μm^2^. Volume is assumed as 500×500×500 μm.

As is evident from the plots, the LS-PSF provides the highest SNR for low excitation intensities. As more power is being used, the SNR saturates due to the finite fluorescence lifetime. Finally, the SNR decreases as the signal is unchanged while the background and noise continues to increase. The best performance of a DL scheme is the one in which a single, DL excitation spot is sparsely scanned over the sample (sparse-PSF; see Fig. 1b and Fig.3, black curve). Furthermore, although the absolute values of the schemes employing a sparsely sampled diffraction-limited PSF (black curve) and LS-PSF (red curve) are comparable, sparse-PSF requires higher intensities while providing no enhancement in spatial resolution compared to LS-PSF.

It is worth noting that although different photo-damage mechanisms are widely debated in the current literature with a range of different thresholds provided, power densities above 20 nJ/μm^2^ have consistently been reported to be detrimental to the viability of cells [50-52], and overall average power limits should be kept below 250 mW to prevent heating of the brain tissue [39]. We have visualized the onset of reported non-linear photodamage by a dashed curve and an average power above 250 mW indicating the regime of possible damage due to tissue heating by a semitransparent curve. In the case of a LS-PSF, average powers close to the heating damage level are reached at lower power densities, as higher average powers are required to excite the enlarged PSF volume. However, for all volume rates shown in Fig. 3, the maximum attainable SNR can be reached using a LS-PSF before heating becomes relevant. This is consistent with our previously reported experimental observations [40] where we imaged such a volume at ^~^3 Hz and did not observe any heat induced immune-reaction. Both LS-PSF and sparse-PSF require higher pulse energies to reach the same average power due the lower repetition rate. We note that the maximum SNR can in general be reached in all investigated scenarios before non-linear photodamage sets in. Moreover, a dwell time <<250 ns is required in order approach voxel rates useful for volumetric imaging on physiological timescales over large volumes. For simplicity, here we have assumed no temporal overhead due to mechanical scanning of the excitation spot, such that the voxel rate is the inverse of the dwell time, i.e. volumes per second, Vps = 1*/Δt.*

In order to investigate the expected volumetric imaging rates that each of the imaging modalities would enable, at each level of power density we looked at the volume rate as a function of the SNR (Fig. 4). The panels show that at any given power density, the sculpted PSF can achieve the highest volume imaging rate with sufficient SNR. Again, the low performance at low volume rates can be attributed to fluorophore saturation and therefore increased background levels compared to the saturated signal.

**Fig. 4:**
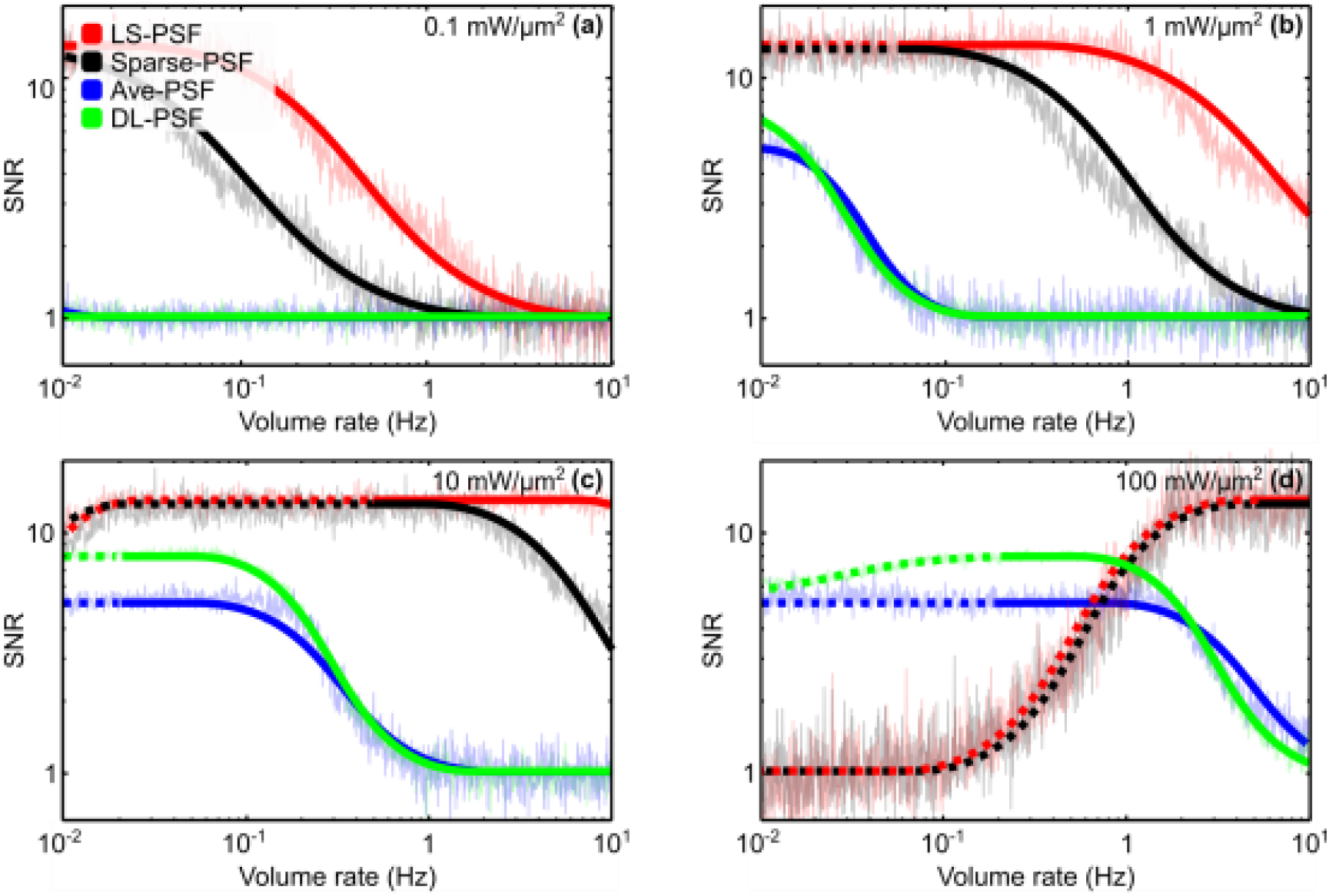
Signal-to-noise ratio as a function of resulting volume recording rate for the different imaging modalities LS-PSF (light sculpted), sparse-PSF (diffraction-limited PSF with sparse scanning), ave-PSF (diffraction-limited PSF and averaging), and DL-PSF (diffraction-limited). The SNR is plotted for (a) 0.1 mW/μm^2^, (b) 1 mW/μm^2^, (c) 10 mW/μm^2^, and (d) 100 mW/μm^2^ incident power density. Dashed lines mark pulse energy densities above the reported onset of photodamage of 20 nJ/μm^2^. Volume is assumed as 500×500×500 μm.

### Performance of various imaging modalities for extraction of neuronal activity

Current research in systems neuroscience is focused on interpreting the neuronal activity of single neurons and neuronal networks. Therefore, any Ca^2+^ imaging modality must produce raw images from which one can manually or through automatic processes, detect single neurons, distinguish them from one another as well as from the background (‘neuropil’), and extract their Ca^2+^-mediated fluorescence intensity modulation (defined as DF/F) over multiple images (i.e. time) in a reliable manner. For our investigation, we chose to benchmark the raw imaging movies from our simulation against a well-known statistical method based on principal and independent component analysis (PCA/ICA) [53]. This method semi-automatically identifies spatial filters, which correspond to individual neurons, and extracts their corresponding fluorescence time traces. Spatial constraints imposed on the ICA segmentation such as size and morphology can further be used to reject non-neuronal signals contributing to the identified spatial filters.

Figure 5 shows examples for the results of the PCA/ICA analysis on our simulated movies obtained with our proposed LS-PSF modality. Fig. 5a and b display the comparison of an extracted Ca^2+^ time trace to the ground truth. The segmentation analysis yields the Ca^2+^ traces of the active neurons as well as their spatial positions. In order to ascertain how well the different imaging modalities are able to correctly identify and resolve single neuronal soma, we compared the identified positions to the ground truth (see Fig. 5c for an example spatial map). Next, we computed Pearson correlation coefficients for the extracted traces with the ground truth dynamics. Figure 5d shows an example of a correlation coefficient histogram. For the following, we regard a correlation coefficient below 0.5 as caused by a false positive in the segmentation analysis.

**Fig. 5:**
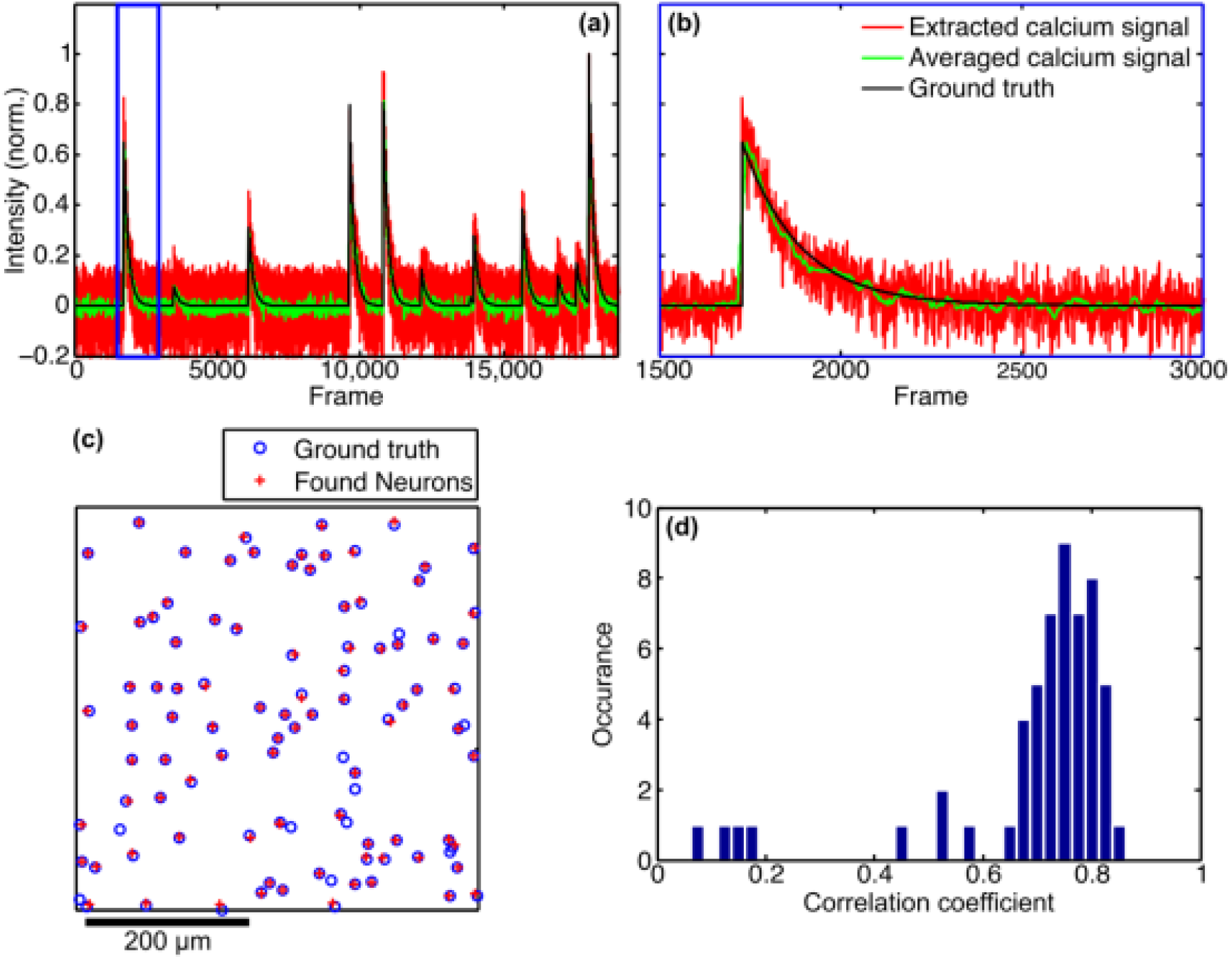
Simulation of Ca^2+^ dynamics obtained with LS-PSF excitation. (a) Example of an extracted calcium signal (red curve: raw signal, green curve: 10 times averaged) and comparison to the ground truth. (b) Zoom-in on the marked (blue box) region in panel a. (c) Example spatial map comparing the identified neurons by the ICA analysis to the ground truth. Here, >90% of neurons are identified. (d) Example histogram of the correlation coefficients for the identified neuron signals with the ground truth dynamics. Correlations <0.5 are regarded as false and likely stem from incorrect neuronal position assignments.

Figure 6 summarizes the performance of the various imaging approaches regarding their ability to correctly identify and resolve single neuronal soma. We note that in this analysis, properties such as the ratio of the PSF to the neuron size are implicitly reflected in the results of the PCA/ICA extracted traces. Therefore, these results also reflect the differences in SNR as they are caused for instance by the different spatial resolution of the four investigated imaging modalities. In this respect as the overall figure of merit of the imaging modalities the percentage of correctly found neuronal signals can be considered. Figure 6a shows the fraction of correctly identified neurons, i.e. the identified neurons subtracted by the false positives. As can be seen the LS-PSF scheme identifies more neurons at lower power densities compared to the alternative, diffraction-limited approaches. We also show in Fig. 6b the identified true neurons as function of the volume rate. The LS-PSF achieves the highest percentage of correctly identified neurons at all volume rates, compared to the other imaging modalities.

**Fig. 6:**
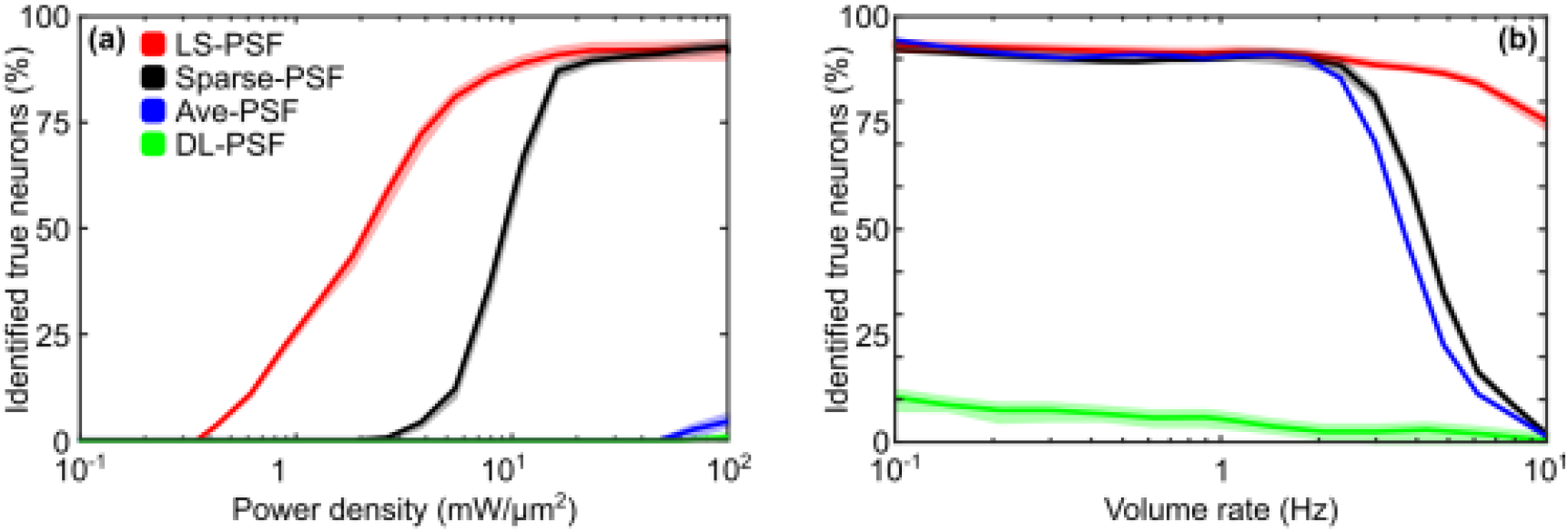
Performance of PCA/ICA analysis on artificial data sets. Identified true neurons as a function of power density for a fixed volume rate of 4 Hz (a), as function of volume rate for a fixed power density of 10 mW/μm^2^ (b). Plotted is the mean +/− std of multiple simulations runs.

## Discussion

In this work, we have investigated new approaches to high-speed volumetric imaging of Ca^2+^ dynamics in scattering tissue, such as the mouse brain. We have quantitatively compared conventional two-photon imaging based on the scanning of a diffraction-limited PSF to other, modified, imaging modalities e.g. with a light-sculpted PSF using numerical simulations. We would like to note that the LS-PSF approach is a holistic optimization where several parameters have been optimized together such as the excitation volume and the number of laser pulses per excitation voxel. While in general an isotropic PSF should always be favorable when imaging a sample volume with no specific axis of anisotropy, we also found that only our LS-PSF scheme has the potential to image volumes whose size (500×500×500 μm) and acquisition speed (>5 Hz) become relevant for current system neuroscience questions where spatial resolution can be traded to some extend for volume size and neuronal recording capacity.

In a recently published experimental work [40], we demonstrated that the LS-PSF is a powerful approach for performing large-scale Ca^2+^ imaging of neuronal activity of large neuronal populations about half of a cortical column, which is often regarded as the computational unit of the brain [54]. The combination of a PSF sculpted in 3D to match the needed resolution of the object of interest (cortical cell bodies) and the one pulse per voxel excitation scheme has allowed us to faithfully extract the activity traces of thousands of neurons distributed in large networks. We expect that similar holistic approaches such as the one presented here will be crucial for optimizing the performances of future imaging systems to record brain dynamics on ever increasing spatial and temporal scales which are essential to gain new insights into the computational principles of information processing and validation of computational models of the mammalian cortex [55-57].

While our investigation and results were obtained with a very specific application in mind, our imaging approach might also find promising applications in other biological fields that rely on multi-photon absorption processes or fluorescence imaging, such as cell and biological population imaging in scattering tissues or more generally in material processing including laser writing, or high-throughput screening. On a more general note, our work also points to a more crucial principle besides the holistic approach to the optical design when developing imaging modalities: Maximizing the use of computational tools in order to relax the constraints and uncertainties on the optics and mechanics, as these usually come at much higher cost. With current efforts conceiving ever more complex imaging modalities, we reckon this topic will become more and more important in future optical engineering efforts.

## Acknowledgements

We thank T. Nöbauer (The Rockefeller University) for datasets to validate our simulations. R.P. acknowledges the Vienna International Postdoctoral Program (VIPS) Program of the Austrian Federal Ministry of Science and Research and the City of Vienna. This work was supported through funding from the US National Institutes of Health (NIH) award 1U01NS094263-01, the Intelligence Advanced Research Projects Activity (IARPA) via Department of Interior/Interior Business Center (DoI/IBC) contract number D16PC00002. The US Government is authorized to reproduce and distribute reprints for Governmental purposes notwithstanding any copyright annotation thereon. The views and conclusions contained herein are those of the authors and should not be interpreted as necessarily representing the official policies or endorsements, either expressed or implied, of IARPA, DoI/IBC, or the US Government.

